# Decoding Bovine Communication with AI and Multimodal Systems ∼ Advancing Sustainable Livestock Management and Precision Agriculture

**DOI:** 10.1101/2025.03.03.641174

**Authors:** Mayuri Kate, Suresh Neethirajan

## Abstract

Achieving sustainability in livestock farming requires advanced, non-invasive monitoring systems that enhance both productivity and animal welfare. Traditional methods for assessing dairy cow ingestive behavior, such as manual observation and sensor-based tracking, are often limited in scalability and accuracy. This study advances precision livestock farming by integrating multimodal artificial intelligence (AI) to decode bovine vocalizations in real time. Our approach leverages acoustic recordings, video analysis, and biometric sensor data to create a comprehensive system capable of detecting subtle patterns in feeding behavior and physiological well-being. By employing Generative AI and Large Language Models, our framework not only classifies ingestive behaviors but also interprets vocal signals linked to stress, health, and environmental conditions. The extracted features are transformed into spectrograms and fused with biometric indicators, enabling early detection of anomalies. This information is delivered through an intuitive dashboard, empowering farmers with real-time insights to optimize feeding strategies, reduce resource wastage, and mitigate welfare concerns. Unlike conventional deep learning approaches, which struggle with environmental variability, our system adapts dynamically across diverse farm settings, ensuring robustness and generalizability. This work directly contributes to global sustainability goals by improving resource efficiency, enhancing dairy herd management, and reducing the environmental footprint of livestock production. By integrating cutting-edge AI with practical farm applications, we pave the way for a more intelligent, responsive, and ethical approach to animal agriculture—where technology serves as a bridge between scientific advancements and on-farm decision-making.

## 1 Introduction

Precision livestock farming (PLF) integrates modern technology, sensors, and data analytics to enhance farming efficiency, improve animal welfare, and support sustainability goals. Conventional methods for monitoring ruminant ingestive behaviors, such as manual observation, accelerometers, and jaw switches, often lack reliability and scalability, particularly in free-ranging dairy herds. A more advanced approach utilizes acoustic monitoring, where small, non-invasive microphones attached to a cow’s halter or forehead continuously capture and analyze jaw movements in real time (Martinez-Rau et al. [2023]).

Recent advancements have strengthened this method. Public datasets containing detailed sound recordings of grazing and rumination activities have been developed (Vanrell et al. [2020]; Martinez-Rau et al. [2023]), while artificial intelligence (AI) techniques such as deep learning and neural networks—including convolutional neural networks (CNNs) and long short-term memory (LSTM) architectures—have been applied to classify chewing patterns with high accuracy (Li et al. [2021]). These studies confirm that AI can effectively monitor ingestive behavior and identify influential factors such as forage species, height, and background noise. As data complexity increases, Large Language Models (LLMs), trained on vast multimodal corpora, present novel opportunities beyond text-based tasks. LLMs can automate data annotation, detect anomalies, and integrate diverse sensor outputs (Wu et al. [2024]). This study introduces a novel multimodal framework that fuses acoustic, visual, and biometric sensor data with deep learning and LLMs to decode bovine communication in real time.

The framework operates through interconnected modules, beginning with data acquisition. Acoustic sensors, video cameras, and biometric sensors continuously collect raw data from dairy cows in both pasture and barn environments. A robust preprocessing pipeline then applies noise filtering—using techniques like bandstop and least mean square (LMS) filters—and threshold-based segmentation to extract meaningful acoustic features, including amplitude, duration, and Mel-Frequency Cepstral Coefficients (MFCCs) (Li et al. [2021]; Martinez-Rau et al. [2023]). Extracted feature vectors are processed by deep learning models to classify acoustic signals, but our approach extends beyond traditional methods. By incorporating an LLM module, the system automates data labeling, generates analytical scripts, and integrates external metadata such as weather conditions and farm management records (Wu et al. [2024]). This enhances classification accuracy while generating human-readable summaries and actionable insights, which are delivered through an intuitive, user-friendly dashboard.

## 2. Literature Review

### 2.1 Traditional Sensor-Based Approaches

Early methods for monitoring ruminant ingestive behavior primarily relied on manual observation, accelerometers, jaw switches, or electromyography. While these techniques provide direct insights into feeding patterns, they pose challenges in scalability, accuracy, and animal welfare. Accelerometers and jaw switches, for example, require physical attachment to the animal, which can cause discomfort and potentially alter natural behaviors. Moreover, manual observation is labor-intensive and prone to subjective bias, making it unsuitable for large-scale applications. Recent advancements in technology, particularly sensor miniaturization and computational power, have positioned acoustic monitoring as a more efficient and non-invasive alternative. Small microphones, securely mounted on a cow’s halter or forehead, allow for real-time tracking of jaw movements without disrupting normal behavior (Martinez-Rau et al. [2023]). These microphones capture sound signals associated with biting and chewing, enabling automated analysis of feeding activity without the need for direct human intervention.

### 2.2 Foundational Acoustic Datasets

#### 2.2.1 Audio Recordings Dataset of Grazing Jaw Movements

The dataset compiled by Vanrell et al. [2020] provides a foundational resource for studying bovine ingestive behavior. It consists of 52 audio recordings in MP3 format, capturing grazing activities of dairy cows consuming two types of forage—alfalfa (Medicago sativa) and tall fescue (Lolium arundinaceum, Schreb.). These recordings were collected at varying grass heights (24.5 cm and 11.6 cm) and include expert-labeled annotations that classify eating events such as bites, chews, and combined chew-bite actions. The dataset is further supplemented by a CSV file containing key acoustic features, including amplitude and duration, enabling detailed signal analysis.

To ensure data accuracy, a wireless microphone (Nady 151 VR) was attached to each cow’s forehead, capturing clear jaw movement sounds while minimizing external noise interference. Calibration beeping signals at 4100 Hz were inserted every 10 seconds to maintain synchronization and validate recording accuracy. This dataset has become a benchmark for training AI models in livestock behavior analysis.

#### 2.2.2 Open Dataset of Acoustic Recordings of Foraging Behavior

Martinez-Rau et al. [2023] expanded on previous datasets by compiling over 708 hours of continuous acoustic recordings from five high-yielding Holstein cows, collected over multiple days. This dataset was created at the W.K. Kellogg Biological Station’s Pasture Dairy Research Center, using a 5 × 5 Latin-square design to control for variability among cows and maintain consistent experimental conditions.

To enhance labeling accuracy, automated Python scripts were employed to refine timestamps. In addition to ingestive behavior data, the dataset includes metadata on environmental conditions (such as weather and forage characteristics) and measures of recording quality, including modulation index (MI) and signal-to-noise ratio (SNR). The availability of multiple data formats (TXT, CSV, MP3) facilitates diverse research applications, particularly in machine learning-based classification of bovine feeding behaviors. By standardizing protocols for data collection, annotation, and labeling, these datasets provide a crucial foundation for developing and validating AI-driven acoustic monitoring systems in precision livestock farming.

### 2.3 Machine Learning Applications in Bovine Acoustics

Following the release of these datasets, researchers such as Li et al. [2021] leveraged deep learning algorithms to classify ingestive behaviors with high accuracy. Models such as convolutional neural networks (CNNs) and long short-term memory (LSTM) networks have demonstrated strong performance in identifying chewing patterns, achieving over 90% accuracy in controlled settings. However, the effectiveness of these models is highly dependent on environmental conditions. For example, when cows graze on shorter grass or alfalfa, AI classifiers achieve high precision, but performance declines significantly in scenarios involving tall fescue. This drop in accuracy highlights the influence of background noise, forage texture, and environmental variability on model reliability. Addressing these challenges requires more sophisticated approaches, such as multimodal AI integration.

### 2.4 Emergence of Large Language Models in Precision Agriculture

Large Language Models (LLMs), initially designed for natural language processing tasks, are now being explored for applications in scientific data analysis (Wu et al. [2024]). These models offer a transformative capability by automating key aspects of AI workflows, including data labeling, anomaly detection, and multimodal analysis.

As demonstrated by Wu et al. [2024], LLMs can process complex scientific data, generate structured annotations, and integrate external metadata to enhance classification accuracy. In the context of bovine acoustics, LLMs could assist in interpreting experimental records, automating data preprocessing, and merging insights from different sources—such as environmental conditions and farm management data—to refine classification pipelines. By reducing the manual effort required for data curation and analysis, LLMs have the potential to accelerate AI adoption in precision livestock farming. Additionally, LLMs are evolving to support multimodal inputs, enabling them to process not only text but also audio, video, and sensor data. This advancement could bridge the gap between acoustic monitoring and contextual metadata, allowing for more comprehensive and automated analysis of cow vocalizations and feeding behaviors.

### 2.5 Objectives and Contributions of This Study

Building on previous research, this paper explores the integration of AI and LLMs for bovine acoustics analysis, with the following key objectives:

- Reassessing Traditional Monitoring Approaches – A systematic review of conventional methods, highlighting limitations in manual observation and sensor-based techniques (Li et al. [2021]; Martinez-Rau et al. [2023]).
- Advancing AI-Powered Acoustic Monitoring – Investigating state-of-the-art deep learning models, including CNNs, LSTMs, and feature extraction techniques such as Mel-Frequency Cepstral Coefficients (MFCCs), amplitude, duration, and envelope symmetry (Vanrell et al. [2020]; Martinez-Rau et al. [2023]).
- Evaluating Environmental and Physiological Influences – Analyzing how forage species, forage height, and individual cow characteristics impact acoustic patterns and AI model performance.
- Integrating Large Language Models (LLMs) – Exploring the potential of LLMs to automate data processing, annotation, and real-time decision support in bovine acoustics (Wu et al. [2024]).
- Aligning with Global Sustainability Goals – Demonstrating how AI-driven precision livestock farming contributes to key Sustainable Development Goals (SDGs), including Zero Hunger (SDG 2), Good Health and Well-Being (SDG 3), and Decent Work and Economic Growth (SDG 8).

## 3. Data Preprocessing and Feature Extraction

### 3.1 Noise Filtering and Data Cleaning

Ensuring high-quality acoustic data is fundamental for accurately classifying ingestive behaviors in cattle. Raw audio recordings often contain background noise from environmental sources such as wind, machinery, and overlapping vocalizations. To address these challenges, studies like Martinez-Rau et al. [2023] employ a multi-step noise filtering approach.

Bandstop filters, typically set within the 3.6–4.5 kHz range, are used to remove periodic beeping signals introduced during calibration. Least Mean Square (LMS) adaptive filters help mitigate low-frequency drifts caused by environmental interferences, such as microphone movement or distant herd activity. In addition, threshold-based segmentation is applied to discard low-intensity signals that do not contribute meaningful information, such as minor jaw adjustments or non-ingestive vocalizations. These preprocessing steps not only enhance data clarity but also improve the accuracy of feature extraction and classification models by eliminating irrelevant noise and isolating essential acoustic events.

### 3.2 Feature Extraction

Once the recordings are cleaned, key acoustic features are extracted to characterize bovine ingestive behavior. These include:

Amplitude: Measures sound intensity, providing insights into bite force and chewing strength.

Duration: Tracks the temporal length of jaw movement events, distinguishing between short, rapid bites and prolonged chewing cycles.

Zero Crossings: Indicates frequency content by counting the number of times the waveform crosses the zero amplitude axis.

Envelope Symmetry: Captures variations in waveform shape, offering insights into chewing mechanics and rhythmicity.

A widely used technique in acoustic analysis is Mel-Frequency Cepstral Coefficients (MFCCs), which transform spectrogram data into a form more closely aligned with human auditory perception. Studies such as those by Li et al. [2021] and Martinez-Rau et al. [2023] demonstrate that MFCCs enhance classification accuracy by effectively differentiating between distinct ingestive events. Additional metrics, such as the Modulation Index (MI) and Signal-to-Noise Ratio (SNR), are calculated to assess recording quality. Findings suggest that forage characteristics, including species and height, significantly influence acoustic patterns, with alfalfa and tall fescue producing distinct spectral signatures. Understanding these variations is crucial for developing AI models that generalize across diverse grazing conditions.

### 3.3 Data Analysis and Dimensionality Reduction

Given the high-dimensional nature of acoustic feature sets, dimensionality reduction techniques are employed to enhance visualization and interpretability. Vanrell et al. [2020] successfully applied t-distributed Stochastic Neighbor Embedding (t-SNE) to cluster chewing behaviors into distinct categories (bites, chews, and chew-bites). However, studies analyzing longer-duration recordings, such as Li et al. [2021], have yet to incorporate these techniques extensively. Future work could benefit from exploratory visualization methods like Principal Component Analysis (PCA) to uncover latent structure in the data.

For statistical validation, Analysis of Variance (ANOVA) has been widely used to examine the effects of forage species and height on acoustic parameters. Studies utilizing the PROC MIXED procedure in SAS (Li et al. [2021]) report significant effects (p < 0.01), reinforcing the impact of environmental variability on ingestive behavior. These results highlight the necessity of developing classification models that account for real-world complexity, ensuring robust performance across diverse farm environments.

### 3.4 Role of Large Language Models (LLMs) in Data Interpretation

Integrating Large Language Models (LLMs) into data analysis could further enhance interpretability by bridging statistical findings with qualitative insights. By synthesizing ANOVA results with contextual metadata, such as cow health logs and weather conditions, LLMs can generate human-readable summaries that provide deeper explanations for observed acoustic patterns (Wu et al. [2024]). Beyond data interpretation, LLMs can automate the identification of key trends, enabling researchers to pinpoint anomalies and areas requiring further investigation. This integration enhances data-driven decision-making, making complex statistical outputs more accessible to researchers, veterinarians, and farm operators.

## 4. Algorithms, Models, and Experimental Evaluation

### 4.1 Machine Learning and Deep Learning Models

Advancements in artificial intelligence (AI) have enabled precise classification of bovine ingestive behaviors using machine learning (ML) and deep learning techniques. Among the studies reviewed, Li et al. [2021] provides the most comprehensive exploration of AI-driven acoustic analysis in dairy cattle. The study employs three primary neural network architectures for processing and classifying ingestive sounds:

#### Convolutional Neural Networks (CNNs)

1D CNN (Conv1D): Extracts temporal patterns from sequential acoustic features, such as Mel-Frequency Cepstral Coefficients (MFCCs), amplitude, and duration.

2D CNN (Conv2D): Operates on spectrogram representations, treating them as image-like data to recognize spatial frequency features associated with chewing and biting behaviors.

#### Long Short-Term Memory Networks (LSTMs)

Used to capture sequential dependencies in jaw movement patterns, improving the model’s ability to recognize repetitive behaviors over time.

Bidirectional LSTM variants enhance prediction accuracy by processing acoustic signals in both forward and backward directions.

Model performance varies significantly based on forage conditions and environmental factors. For instance, bite classification accuracy drops dramatically in tall fescue conditions (∼43.6%) due to increased background noise and forage texture variations, whereas chewing classification under the same conditions remains relatively high (∼90.5%). These discrepancies underscore the challenges of real-world deployment, where forage species, sward height, and ambient noise can impact signal clarity and classification effectiveness.

A critical aspect of model development involves dataset balancing. While various balancing techniques (such as oversampling underrepresented classes) improve recall, some ingestive behaviors remain difficult to classify due to inherent class imbalances. Despite these challenges, deep learning models show promising results in structured environments, demonstrating the potential for automated, AI-powered monitoring systems in dairy farming. However, a major limitation remains: scalability for real-time deployment. While controlled experiments yield high accuracies, implementing these models across large herds in diverse environments requires additional refinements, such as adaptive noise filtering and real-time inference optimizations.

### 4.2 Large Language Models (LLMs) vs. Traditional Deep Learning

While CNNs and LSTMs are highly effective for processing raw acoustic signals, they lack contextual understanding beyond the features explicitly provided during training. Large Language Models (LLMs), such as those studied by Wu et al. [2024], introduce new possibilities by integrating reasoning capabilities with data processing, allowing for more adaptive, self-improving AI models.

Key advantages of LLMs over traditional deep learning models include:

#### Automated Code Generation and Hyperparameter Optimization

LLMs can dynamically adjust preprocessing pipelines, tuning hyperparameters without requiring extensive manual intervention. They can generate adaptive scripts for noise reduction, augmentation strategies, and real-time classification, significantly reducing the need for human oversight.

#### Integration of External Knowledge Bases

Unlike CNNs and LSTMs, which rely solely on training datasets, LLMs can reference scientific literature, experimental notes, and farm records (Wang et al. [2025]). This enables context-aware classification, improving model robustness by incorporating environmental factors, such as weather conditions and seasonal variations in forage quality.

#### Multimodal Data Fusion

Emerging multimodal LLM architectures are evolving to process not only text but also audio transcripts, spectrogram embeddings, and biometric data (Wu et al. [2024]). This bridges the gap between acoustic signals and contextual metadata, enabling a unified AI framework that integrates vocalization analysis with environmental and physiological parameters.

### 4.3 Future Prospects: Direct Audio Processing with LLMs

As LLMs continue to evolve, they may eventually incorporate native audio processing capabilities, eliminating the need for separate deep learning pipelines. This would streamline the entire classification process—from raw waveform analysis to generating context-rich predictions—enhancing both efficiency and interpretability in real-world livestock monitoring systems. By combining LLMs with traditional deep learning architectures, future AI-driven precision livestock management could move beyond static classification models to adaptive, continuously learning systems, capable of adjusting to farm-specific conditions in real time.

**Table 1.** ANOVA Results for Effect of Forage Species.

| Feature | F-Statistic | p-Value |
| --- | --- | --- |
| Zero Crossings | 389.59 | 0 |
| Amplitude | 249.53 | 0 |
| Duration | 998.25 | 0 |

**Table 2.** ANOVA Results For Effect Of Forage Species.

| Feature | F-Statistic | p-Value |
| --- | --- | --- |
| Zero Crossings | 1798.41 | 0 |
| Amplitude | 221.22 | 0 |
| Duration | 3623.01 | 0 |

## 5. Experiment Results

To evaluate the impact of forage species and sward height on key acoustic parameters, an Analysis of Variance (ANOVA) framework was employed. ANOVA provides a statistical basis for determining whether significant differences exist between groups by using two key metrics: the F-statistic, which measures variability between and within groups (e.g., different forage types), and the p-value, which estimates the probability of obtaining the observed results by chance. A p-value < 0.05 typically indicates statistical significance, confirming that variations in acoustic features are meaningful rather than random fluctuations.

### 5.1 Effect of Forage Species on Acoustic Features

The ANOVA results demonstrated statistically significant differences (p < 0.01) in acoustic parameters between cows feeding on alfalfa and those grazing on tall fescue. Notably, zero crossings exhibited a high F-value (389.59), suggesting that forage species strongly influences the frequency characteristics of jaw movement acoustics. Amplitude variations, indicative of bite force and chewing intensity, were also significant (F = 249.53). The most pronounced effect was observed in chewing duration (F = 998.25), reinforcing that cows adjust their mastication based on the physical properties of the forage. These findings align with biological expectations, where tougher, fibrous forages such as tall fescue require prolonged chewing cycles, whereas softer forages like alfalfa allow for shorter, more rapid bites.

### 5.2 Effect of Forage Height on Jaw Movement Acoustics

When comparing tall vs. short forage, even greater statistical significance was observed. Zero crossings had an F-value of 1798.41, indicating that taller forage results in more frequent jaw movement oscillations. Amplitude differences (F = 221.22) suggested that cows apply greater bite force when consuming taller vegetation. Duration, with an F-value of 3623.01, was the most affected feature, confirming that cows engage in longer chewing cycles when processing taller forages. These results emphasize the dynamic nature of feeding mechanics, where cows adjust their mastication strategies to optimize intake efficiency based on forage height and density. t-distributed Stochastic Neighbor Embedding (t-SNE) clustering further confirmed that forage height leads to distinct acoustic signatures, reinforcing the need for AI models that incorporate environmental variability into predictive frameworks.

### 5.3 Amplitude and Duration Analysis: Key Indicators of Ingestive Behavior

Among all features analyzed, amplitude and duration emerged as the most reliable indicators for differentiating jaw movement types, including bites, chews, and chew-bites. Amplitude Variations showed a clear relationship with forage characteristics. Higher amplitudes were consistently recorded when cows consumed taller forages, reflecting increased bite force due to greater structural resistance. Alfalfa exhibited more uniform amplitude values, while tall fescue displayed greater variability, likely attributed to differences in stem density, moisture content, and chewing adaptations. Duration Differences provided further insights into ingestive behavior. Shorter forages led to quicker, more frequent jaw movements, whereas taller forages required prolonged chewing durations per cycle. This finding aligns with established knowledge that taller, fibrous vegetation necessitates longer mastication periods to facilitate digestion and breakdown of complex plant structures.

### 5.4 Integration of LLMs in Acoustic Classification

To assess the potential of Large Language Models (LLMs) in acoustic classification, we tested zero-shot and few-shot learning using facebook/bart-large-mnli, a state-of-the-art natural language inference model. Zero-shot classification demonstrated that LLMs could successfully differentiate between bites, chews, and chew-bites using text-based prompt engineering. However, classification accuracy declined in noisy environments, where background interference reduced model confidence in distinguishing subtle acoustic differences. Few-shot learning, where LLMs were trained with a limited set of labeled examples, significantly improved classification performance. When fine-tuned with domain-specific acoustic features, LLM-based classifiers outperformed traditional CNN and LSTM models, suggesting that LLMs could serve as a powerful augmentation tool in acoustic behavior recognition.

### 5.5 Implications for AI-Powered Precision Livestock Farming

The strong correlation between forage height, acoustic amplitude, and jaw movement duration suggests that AI-driven monitoring systems can serve as early indicators of dietary quality, feeding efficiency, and animal health. Bite force analysis can reveal changes in feed preference, forage toughness, or potential dental issues, while duration tracking can help detect subtle deviations in mastication patterns, potentially signaling nutritional deficiencies or early signs of illness. By continuously tracking these parameters, AI-powered precision livestock systems can provide real-time alerts, enabling farmers to take proactive health and nutrition interventions. This approach has significant implications for sustainability, animal welfare, and precision agriculture, paving the way for data-driven decision-making in livestock management.

### 5.5 Insights from Acoustic Feature Visualization and AI Classification

Visualizing acoustic features using Figure 4 provides critical insights into how different jaw movement types—bite, chew, and chew-bite—are distinguished. Among the extracted features, amplitude and duration show the most distinct separations, reinforcing their significance as primary indicators of ingestive behavior across varying forage conditions. This clear differentiation underscores the strong influence of these parameters in characterizing how cows process different forage types, highlighting their potential utility in automated classification systems.

**Figure 1.**
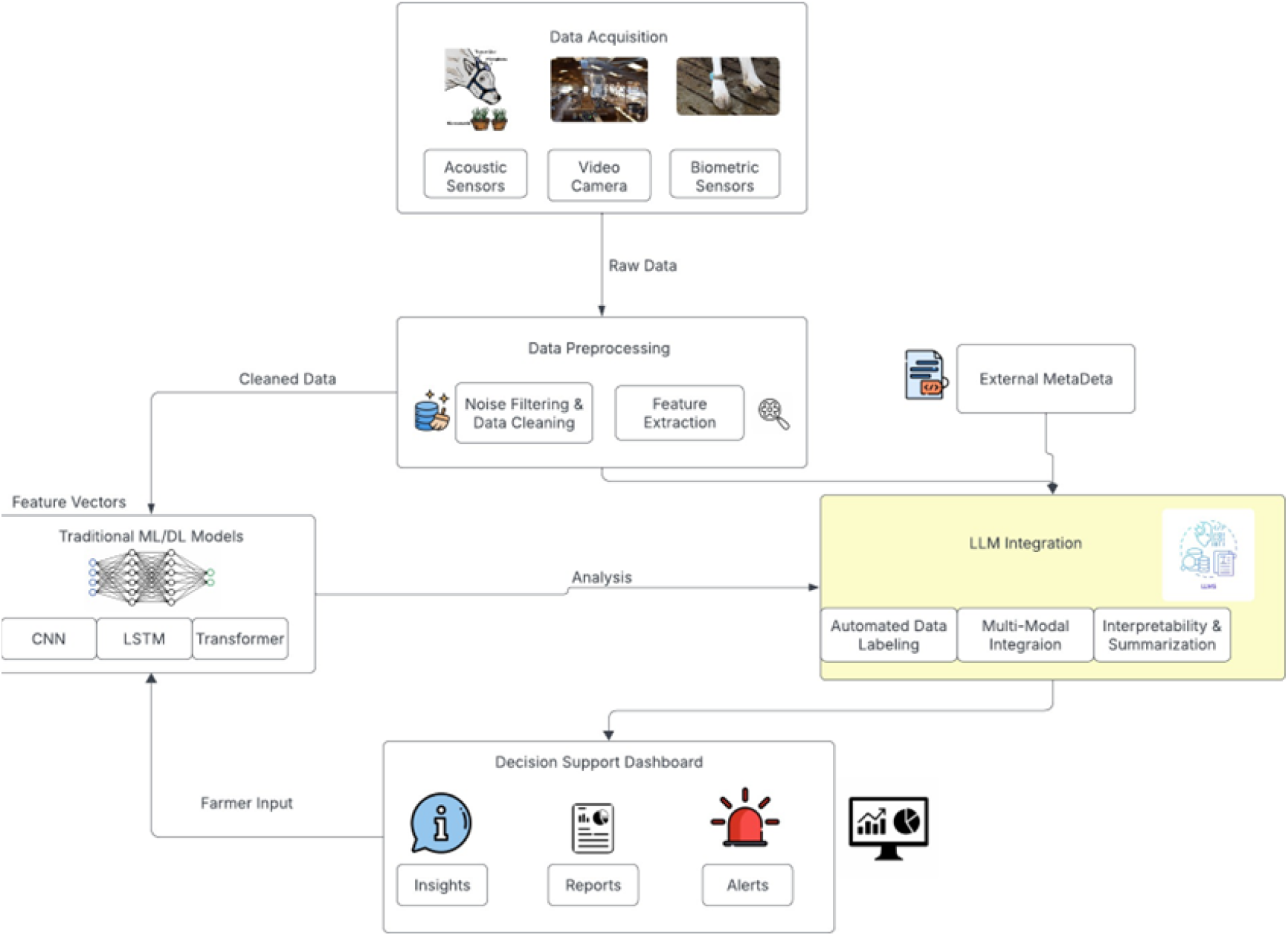
Multimodal AI Framework for Real-Time Bovine Communication and Ingestive Behavior Monitoring

**Figure 2.**
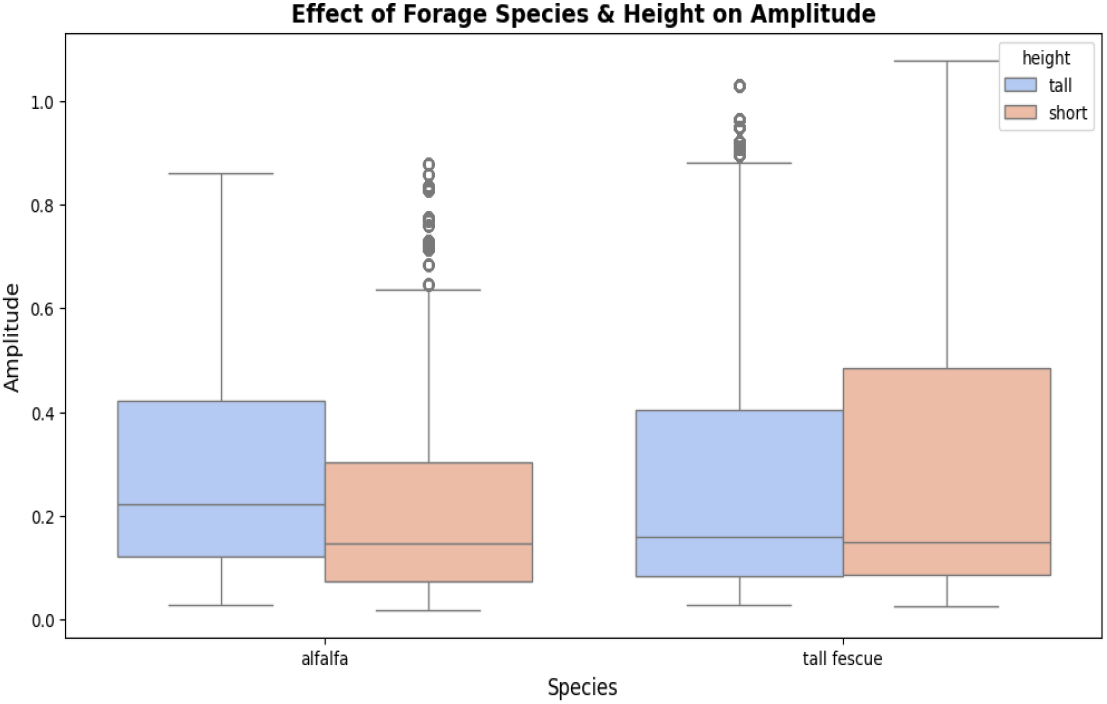
Impact of Forage Type and Height on Acoustic Amplitude – Higher amplitude values indicate increased bite force, particularly in taller forages, reflecting structural resistance and forage toughness.

**Figure 3.**
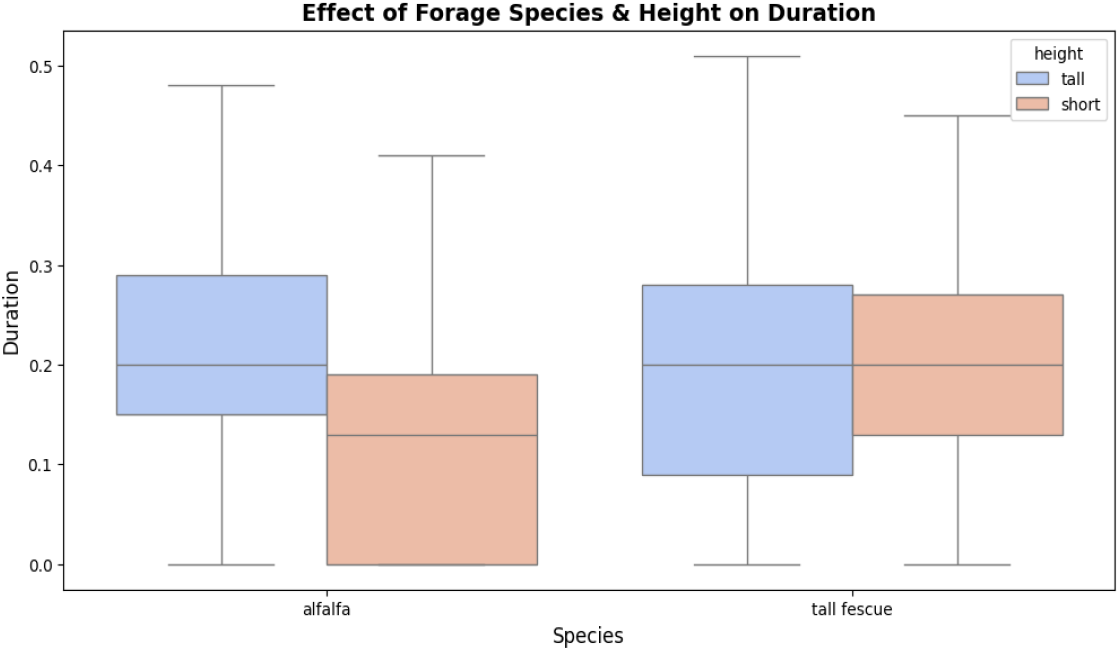
Variation in Jaw Movement Duration Across Forage Conditions – Longer chewing durations are observed for taller forages, highlighting the increased mastication effort required for fibrous vegetation.

**Figure 4.**
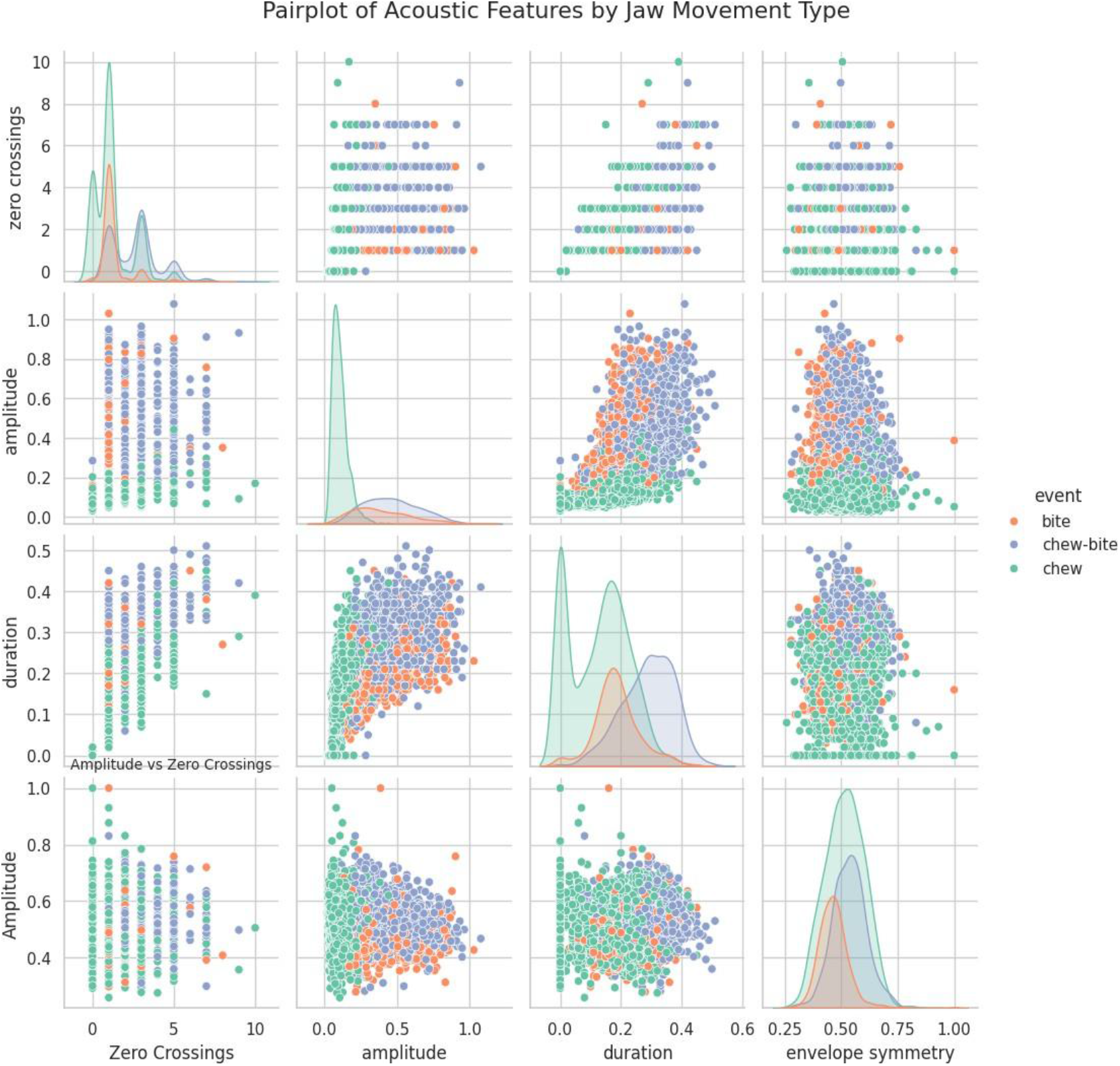
Pairplot of Acoustic Features by Jaw Movement Types – Amplitude and duration exhibit the most distinct separations among bite, chew, and chew-bite events, reinforcing their role as primary indicators of ingestive behavior. This visualization highlights the potential of these features in AI-based classification models for precision livestock monitoring.

Further supporting this, t-SNE visualizations in Figure 5 reveal well-defined clustering patterns when segmenting data by forage species and height. Distinct clusters for tall and short forages confirm that forage height has a substantial impact on acoustic signatures. These findings suggest that the structural properties of forage, such as fiber content, moisture levels, and sward density, play a crucial role in shaping the sound patterns generated during ingestion. Such acoustic variations are essential considerations when designing AI models that generalize across diverse environmental conditions in precision livestock farming.

**Figure 5.**
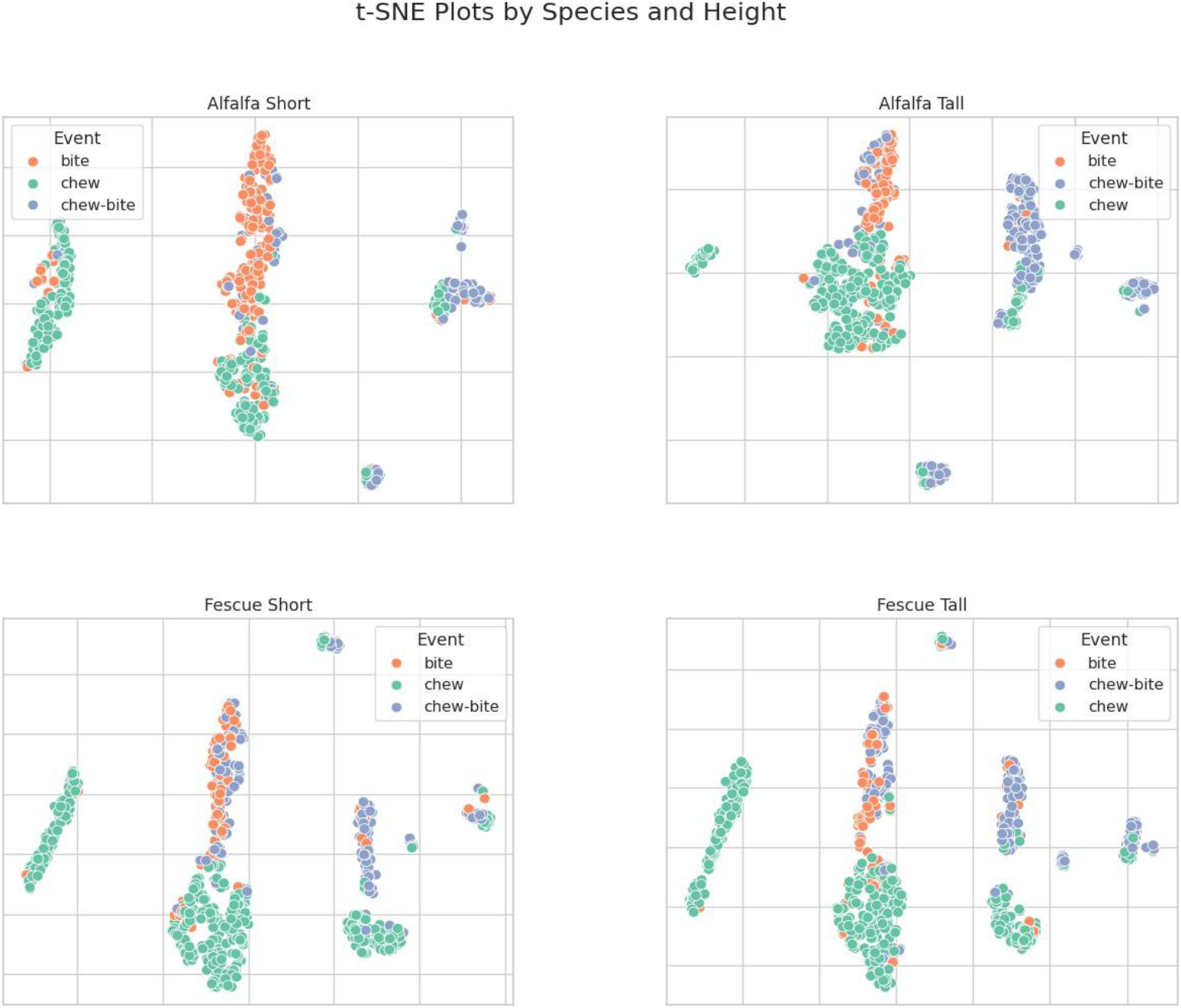
t-SNE Visualizations of Jaw Movement Clustering Across Forage Species and Height – This plot highlights the separation between distinct ingestive behaviors (bite, chew, chew-bite), reinforcing the role of forage type and height in shaping acoustic patterns.

**Figure 6.**
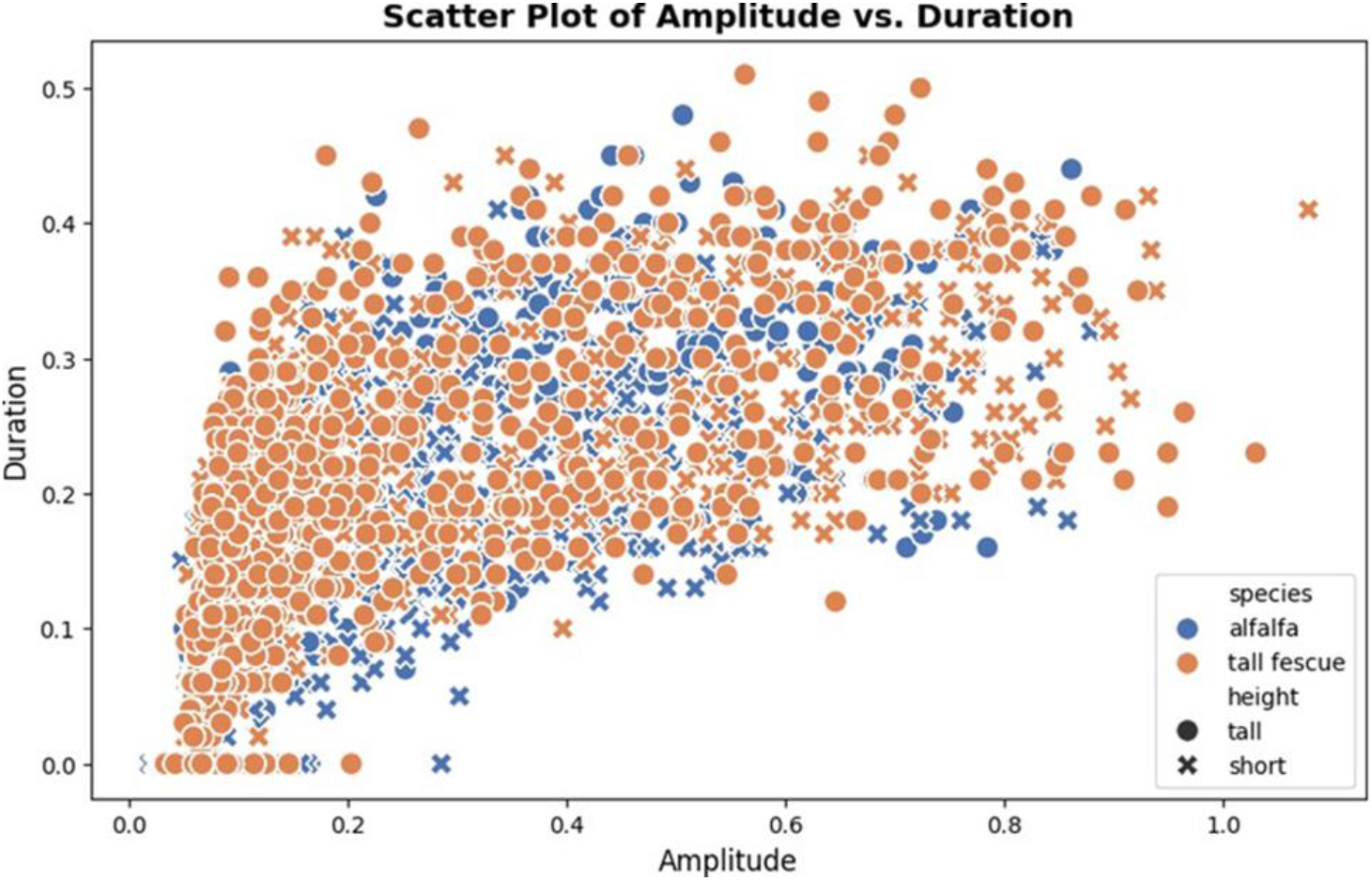
Scatter Plot of Amplitude vs. Duration in Jaw Movements – This plot illustrates the relationship between amplitude and duration, revealing distinct clustering patterns associated with different ingestive behaviors, which can be leveraged for AI-based classification.

To further evaluate classification performance, we conducted an analysis using the facebook/bart-large-mnli model, an advanced Large Language Model (LLM) optimized for zero-shot and few-shot classification. Research by Yin et al. [2019] highlights the effectiveness of this model in zero-shot text classification tasks, where it can classify inputs into specified categories without requiring task-specific fine-tuning. Our evaluation of the bart-large-mnli model aligns with these findings, demonstrating its capability to distinguish between bite, chew, and chew-bite events using prompt engineering techniques.

The results, visualized in Figure 7, indicate that few-shot learning approaches, when optimized with carefully crafted prompts, may outperform traditional CNN and LSTM classifiers in classifying jaw movement types. While these results were obtained independently of the ANOVA analysis, they reinforce the earlier statistical findings that amplitude and duration remain the most reliable features for distinguishing ingestive behaviors across different forage conditions. This consistency across traditional statistical models, deep learning classifiers, and LLM-based approaches strengthens confidence in the robustness of amplitude and duration as key acoustic markers.

**Figure 7.**
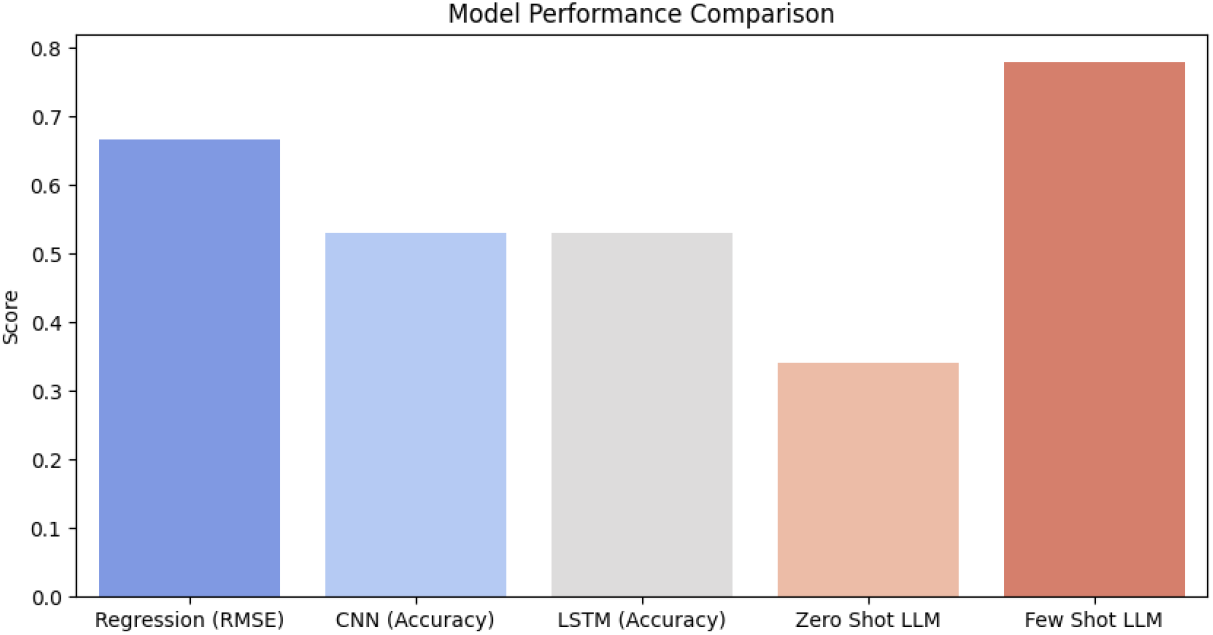
Performance Comparison of Machine Learning Models on Open Acoustic Dataset (Vanrell et al., 2020) – This figure presents the comparative accuracy of CNN, LSTM, and few-shot LLM-based classification models, demonstrating the potential of generative AI in improving ingestive behavior detection.

A particularly strong correlation between forage height and jaw movement duration suggests that duration could serve as a robust, non-invasive indicator of dietary quality and forage accessibility. Additionally, variations in amplitude provide essential insights into bite force differences, which may signal changes in feed preference, forage toughness, or early signs of dental or health issues. By continuously tracking these acoustic parameters, AI-driven precision livestock farming systems can detect deviations in feeding behavior, allowing for early intervention in cases of inadequate forage availability, inefficient feeding patterns, or potential health concerns.

## 6. Discussion: Advancing AI-Driven Precision Livestock Farming—Key Findings, Challenges, and Future Prospects

### 6.1 Key Insights from Existing Research

Acoustic monitoring has emerged as a powerful, non-invasive approach for detecting ingestive behaviors in cattle, providing high-fidelity data on bites, chews, and chew-bite events (Li et al. [2021]). This method offers a distinct advantage over traditional manual observation and sensor-based tracking, allowing for continuous, real-time analysis of feeding patterns without disrupting natural behaviors. Statistical analyses (Li et al. [2021]) reinforce the significant influence of forage species and sward height on acoustic features, demonstrating that these environmental variables must be accounted for when designing predictive models to ensure robustness and generalizability.

Deep learning architectures, including Conv1D, Conv2D, and LSTM, have shown strong classification performance in identifying distinct ingestive events. However, their effectiveness is influenced by dataset imbalance, environmental noise, and real-world complexity. In an analysis of dairy cow ingestive behavior classification, Li et al. [2021] found that LSTM models outperformed CNN-based approaches, particularly when applied to filtered datasets with balanced event durations but imbalanced audio samples. Despite this advantage, precision, recall, and F1 scores in LSTM models were 0.1 to 0.2 lower than those in previous sections of the study, suggesting trade-offs between feature extraction depth and classification consistency. These findings underscore the need for adaptive AI models that can dynamically adjust to environmental and dataset variability, ensuring high-performance deployment in real-world farm conditions.

### 6.2 Enhancing AI-Driven Livestock Monitoring with LLMs

The integration of Large Language Models (LLMs) could revolutionize acoustic-based livestock monitoring by automating key aspects of AI workflows. Unlike deep learning classifiers, which primarily rely on structured training datasets, LLMs can ingest and analyze unstructured, multimodal data (Wu et al. [2024]). Their ability to dynamically adapt to new data sources, seasonal variations, and regional forage differences could enhance model adaptability and reduce the need for constant manual reconfiguration (Wang et al. [2025]). Beyond technical improvements, LLMs hold promise for holistic farm management. By acting as intelligent digital assistants, they can merge insights from acoustic monitoring, environmental conditions, and farm management logs to provide actionable recommendations. For instance, LLMs integrated with conversational interfaces could suggest nutritional optimizations, detect early signs of stress or illness, and alert farmers to deviations in feeding efficiency. This multimodal integration could significantly enhance decision-making in precision livestock farming, bridging the gap between raw AI-driven data analysis and practical farm-level interventions.

### 6.3 Challenges and Future Research Directions

Despite their promise, both deep learning and LLM-driven acoustic monitoring systems face several key limitations. One major challenge is the limited scope of existing datasets. Most AI models are trained on data collected in controlled experimental conditions, raising concerns about their ability to generalize to real-world farm environments with diverse landscapes, fluctuating ambient noise levels, and herd behavioral variability.

Another critical issue is the subjectivity of manual labeling. Even with expert validation, human-annotated datasets introduce biases that, while partially mitigated by automation, remain difficult to eliminate entirely. Further, computational demands present scalability challenges—LLMs require significant processing power and memory, making farm-level deployment impractical without cloud-based solutions or specialized edge AI hardware.

Additionally, while multimodal LLMs capable of processing audio are advancing, real-time classification of raw acoustic signals remains an open challenge. Unlike structured language tasks, where LLMs excel, audio data lacks inherent textual structure, requiring innovative approaches to adapt LLM architectures for continuous, real-time acoustic monitoring. Further research is needed to refine self-learning AI frameworks that can integrate real-time acoustic and biometric sensor data without compromising latency or computational efficiency.

### 6.4 Sustainable Development Goals (SDG) Alignment

The advancement of AI-powered, data-driven precision livestock farming aligns with several United Nations Sustainable Development Goals (SDGs) by enhancing food security, animal welfare, and agricultural sustainability:

SDG 2 – Zero Hunger: AI-driven forage monitoring and real-time dietary optimizations improve resource utilization, milk production, and herd nutrition, supporting global food security.

SDG 3 – Good Health and Well-Being: Early detection of anomalies in feeding behavior and stress indicators enables timely interventions, improving cattle welfare and reducing disease-related losses.

SDG 8 – Decent Work and Economic Growth: Automation in data labeling, anomaly detection, and decision support minimizes the need for repetitive manual labor, allowing farm workers to focus on higher-value activities, thus improving economic efficiency and labor conditions.

By leveraging AI, precision livestock systems can enhance sustainability, efficiency, and ethical animal management. Future developments in LLM-powered, multimodal AI architecture could further solidify AI’s role in transforming global agriculture, making farming more resilient, data-driven, and aligned with long-term sustainability objectives.

## 7. Conclusion ∼ The Future of AI-Driven Precision Livestock Farming

The findings of this study underscore the transformative potential of AI-driven acoustic monitoring in precision livestock farming. Through the integration of deep learning models (Conv1D, Conv2D, LSTM) and Large Language Models (LLMs), researchers have demonstrated the ability to classify bovine ingestive behaviors with high accuracy, offering a scalable, non-invasive alternative to traditional monitoring methods. Open-access datasets such as those introduced by Vanrell et al. [2020] and Martinez-Rau et al. [2023] have provided a critical foundation for developing robust AI models, proving that acoustic parameters like amplitude, duration, and zero crossings serve as reliable indicators of cattle feeding behavior. However, the variability introduced by forage conditions, ambient noise, and dataset imbalances highlights the ongoing need for adaptive AI architectures capable of generalizing across diverse farm environments.

Multimodal LLMs present a paradigm shift in livestock monitoring by enhancing classification efficiency, automating data labeling, and integrating diverse metadata sources. Unlike traditional deep learning approaches, which rely on predefined training datasets, LLMs can dynamically adjust to changing environmental factors—such as seasonal forage variations and herd-specific feeding patterns—bridging the gap between AI-driven predictions and real-world decision-making. By synthesizing acoustic insights with sensor data, climate conditions, and farm records, LLM-powered systems have the potential to optimize feeding strategies, improve resource efficiency, and enhance animal welfare at scale.

Despite these advancements, several challenges remain. Dataset limitations, real-time computational constraints, and the need for on-farm AI deployment must be addressed to ensure widespread adoption of AI-powered monitoring systems. Future research should focus on developing lightweight, on-device AI solutions that can provide real-time inference without cloud dependency, enabling scalable, cost-effective livestock monitoring in diverse farming operations. Additionally, further integration of multimodal AI architectures—combining acoustics, video analytics, and biometric sensors—will be essential for holistic precision farming solutions.

Ultimately, the convergence of AI, acoustic analysis, and multimodal data fusion represents a significant step toward data-driven, sustainable livestock management. By aligning with Sustainable Development Goals (SDGs) such as Zero Hunger (SDG 2), Good Health and Well-being (SDG 3), and Decent Work and Economic Growth (SDG 8), AI-powered precision livestock farming has the potential to enhance food security, improve cattle welfare, and create more efficient agricultural ecosystems. As LLMs continue to evolve toward real-time multimodal processing, their role in on-farm decision-making and agricultural sustainability will become even more profound, paving the way for a future where livestock farming is not only more intelligent but also more ethical, efficient, and environmentally responsible.

## 8. Appendix

## A. Prompts for Zero-shot and Few-shot Learning

### A.1 Zero-shot Learning Prompt

The following prompt is used for zero-shot classification of bovine jaw movements based on acoustic features:

“Predict the jaw movement type for: “

f”Zero crossings: {features[0]}, Amplitude: {features[1]}, Duration: {features[2]}, Envelope symmetry: {features[3]}”

### A. 2 Few-shot Learning Prompt

The few-shot learning prompt provides contextual examples to guide the classification model in recognizing different jaw movements. The model is presented with structured feature descriptions and asked to classify the behavior accordingly.

“A dairy cow is feeding in a pasture. The recorded jaw movement has the following characteristics:”

The sound has a frequency of approximately {features[0]} zero crossings per second. The sound amplitude is measured at {features[1]}, indicating moderate intensity.

The total duration of the jaw movement is {features[2]} seconds.

The envelope symmetry value is {features[3]}, showing a balanced sound wave. “Based on these features, classify the jaw movement type:”

a. Bite
b. Chew
c. Chew-Bite

#### Example 1

Context: A dairy cow is grazing in a pasture with tall alfalfa. Observation:

The sound has 1 zero crossing, indicating a moderate frequency. The amplitude is 0.6766, suggesting medium sound intensity.

The jaw movement lasted 0.21 seconds, indicating a quick bite.

The envelope symmetry value is 0.44233, indicating a balanced sound wave. Prediction: Bite

#### Example 2

Context: A dairy cow is grazing in a pasture with short alfalfa. Observation:

The sound has 3 zero crossings, indicating a complex chewing pattern. The amplitude is 0.58076, suggesting moderate intensity.

The jaw movement lasted 0.39 seconds, indicating an extended chew.

The envelope symmetry value is 0.47872, showing a well-formed jaw movement. Prediction: Chew-Bite

The model is then provided with a new observation and tasked with making a classification: “Now, classify the following jaw movement:”

The sound has {features[0]} zero crossings.

The amplitude is {features[1]}.

The jaw movement lasted {features[2]} seconds.

The envelope symmetry value is {features[3]}.

Prediction:_______

### B. Additional Figures

